# An updated compendium of *Caenorhabditis elegans* RNA-binding proteins and their regulation

**DOI:** 10.1101/2025.03.31.646443

**Authors:** Felicia Peng, John Isaac Murray

## Abstract

Although gene regulation occurs both transcriptionally and post-transcriptionally, systems-level characterizations of RNA-binding proteins (RBPs) are still lacking compared to transcription factors. RNA-binding proteins have gene expression functions that include regulating RNA splicing, localization, translation, and turnover. Mapping the regulatory networks that they are involved in will thus be critical for a comprehensive understanding of gene regulation during development. We updated the wRBP1.0 compendium of putative *C. elegans* RNA-binding proteins to 928 proteins in WS290 and have catalogued their expression and developmental phenotypes using existing functional genomic resources. Many RBP genes are expressed in a cell type- and developmental stage-specific manner in the embryo, emphasizing that RBPs can contribute to distinct gene expression patterns. In addition, RBPs are highly conserved, and their loss can result in a wide range of developmental defects. This updated compendium will provide a resource for functional studies of RBP regulatory networks in *C. elegans*.

## Introduction

Gene regulation is key to specifying different cell fates during development. Although developmental gene expression is primarily framed in terms of transcription factor (TF) regulatory networks, RNA-binding proteins (RBPs) are also important regulators of gene expression (Tamburino et al. 2013; Gerstberger et al. 2014; Iadevaia and Gerber 2015; Haskell and Zinovyeva 2021). Elucidating the regulatory networks that RBPs are involved in will thus be important for a more comprehensive understanding of gene regulation during development.

RBPs have a wide array of functions, as they can interact with both coding and noncoding RNAs (Gerstberger et al. 2014). Co- and post-transcriptional gene expression functions include regulating RNA splicing, localization, translation, and turnover (Gerstberger et al. 2014). RBPs generally contain RNA-binding domains (RBDs), and these domains are typically responsible for mediating sequence-specific interactions between RBPs and their RNA targets (Auweter et al. 2006). In a *Caenorhabditis elegans* compendium of 887 predicted RBPs (wRBP1.0), RBPs were split into four groups based on the protein domains they contain (Tamburino et al. 2013). Group 1 RBPs are expected to act in a gene- and RNA-specific manner and contain one of eight RBDs (RRM, KH, PUF, CCCH, CCHC, DSRBD, RGG Box, La). Group 2 RBPs likely bind in a gene-specific and RNA non-specific manner with one of four RBDs (helicase, PAZ, PIWI, NTF2) and include proteins involved in small RNA pathways. Group 3 RBPs contain one of three domains that could interact either with DNA, protein or RNA, likely in a gene-specific manner (C2H2 SAM, cold shock). Finally, Group 4 RBPs bind RNA in a non-gene-specific manner, and are characterized by Sm/Lsm domains and conservation to RBPs in other organisms, including ribosomal proteins and tRNA synthetases. That compendium showed more complex regulation of RBPs than the proteome as a whole; this included more TFs bound to RBP promoters, longer 3’UTRs that have more binding sites for miRNAs and other RBPs, and more phosphorylation sites.

A systems-level characterization of RBPs, in a similar manner to what has been more expeditiously carried out for TFs, will be important in understanding how this group of proteins may contribute to developmental gene expression. Of particular interest are Group 1 RBPs, which are thought to regulate genes in a specific manner analogous to the way TFs regulate gene expression. Group 1 RBPs include proteins with RNA Recognition Motif (RRM) or K homology (KH) domains, which are among the most prevalent protein domains in metazoans (Cook et al. 2011). wRBP1.0 contained 115 proteins containing an RRM domain and 33 proteins containing a KH domain (Tamburino et al. 2013).

Here, we update the list of predicted RBPs to a current genome annotation, finding a total of 928 proteins. We catalogued the expression and developmental importance of RBP genes throughout development, largely during embryogenesis, using existing functional genomic resources. We found that many RBP genes are expressed in a cell type- and developmental stage-specific manner, which highlights the potential for RBPs to contribute to cell type-specific gene expression patterns. Furthermore, loss of RBPs can result in a wide range of developmental defects. Group 1 RBP genes are enriched for expression in the germline, though many are also enriched in somatic cell types, such as neurons and muscle. The turnover of Group 1 RBP transcripts may also be highly regulated. Altogether, this updated compendium and analysis of different categories of RBPs provides a resource for functional studies of RBP regulatory networks in *C. elegans* development.

## Results

### Updated compendium of predicted RNA-binding proteins

To expand upon the wRBP1.0 compendium of predicted RBPs, we first merged gene names and removed genes that were dead to align with the *C. elegans* genome build WS290. InterPro protein domains were then retrieved from the WormBase ParaSite database (Howe et al. 2017), and the proteome was searched for each of 17 RBDs as in wRBP1.0 (Tamburino et al. 2013). Our search of the updated *C. elegans* proteome resulted in the addition of six new Group 1 RBPs, eight new Group 2 RBPs, 32 new Group 3 RBPs, and one new Group 4 RBP (Supplemental Table S1). In total, our updated compendium has 928 predicted RBPs (Figure 1A; Supplemental Table S1).

**Figure 1.**
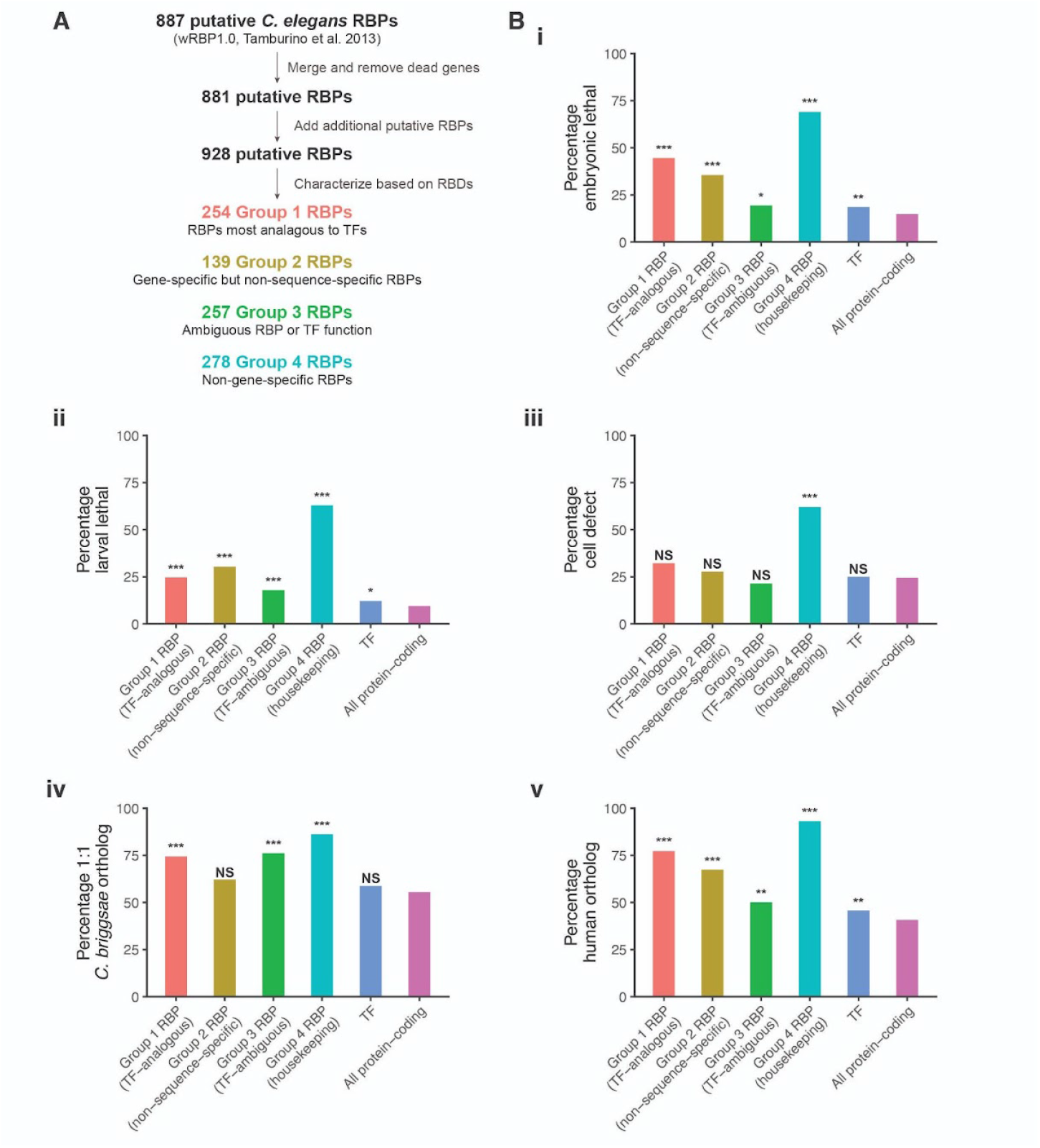
RNA-binding proteins are important for *C. elegans* development. (A) Workflow for updated *C. elegans* RBP predictions. (B) (i, ii, iii) The percentage of genes in each category with an associated developmental defect upon their depletion using RNAi. Data from i and ii came from Wormbase (Lee et al. 2018), and data from iii came from (Xiao et al. 2022). (iv) The percentage of genes in each category with a 1:1 ortholog in *C. briggsae*. Orthology data taken from (Large et al. 2024). (v) The percentage of genes in each category with an ortholog in humans. Orthology data taken from Ortholist 2 (Kim et al. 2018). A Chi-Square Test was used to determine statistically significant differences in each group compared to all protein-coding genes.

For downstream comparative analyses, we limited Group 1 RBPs to those mutually exclusive with TFs (Fuxman Bass et al. 2016); however, proteins predicted to potentially be both RBPs and TFs were categorized as RBPs if they contained a CCCH domain, as several proteins containing this domain have been well-characterized as RBPs that regulate cell fate specification during early *C. elegans* embryogenesis (Elewa et al. 2015). Group 2 RBPs were similarly restricted to non-TFs. As Group 3 RBPs are ambiguous in having RNA- or DNA-binding functions, putative TFs were kept in this group for downstream analyses. Putative RBPs from all four groups were excluded from TFs analyzed. Ultimately, 251 Group 1 RBPs, 135 Group 2 RBPs, 257 Group 3 RBPs, 278 Group 4 RBPs, and 700 TFs were included in our downstream analyses (Supplemental Table S2).

### RNA-binding proteins are important for C. elegans development

To broadly determine the developmental importance of RBPs in *C. elegans*, we examined developmental defects associated with their depletion. RNA interference (RNAi) phenotype data from Wormbase (Lee et al. 2018) revealed that knockdown of 44.6%, 35.6%, 19.5%, and 69.1% of included Group 1-4 RBP genes, respectively, resulted in embryonic lethality (Figure 1B, i; Supplemental Table S1). For most groups this was substantially higher than the percentage of embryonic lethality for TF (18.6%) and all protein-coding (14.9%) genes. Knockdown of 24.7%, 30.4%, 17.9%, and 62.9% of included Group 1-4 RBP genes, respectively, resulted in larval arrest (Figure 1B, ii; Supplemental Table S1). For TF and all protein-coding genes, the percentage of larval arrest was 12.1% and 9.5%, respectively.

Developmental defects upon RNAi knockdown have also been examined in greater detail for over 750 conserved genes in the *C. elegans* embryo using direct cell lineage tracing (Xiao et al. 2022). These conserved genes include 31 Group 1 RBP genes, 18 Group 2 RBP genes, 14 Group 3 RBP genes, 29 Group 4 RBP genes, and 32 TF genes. Depletion of 21.4-62.1% of genes included in the RBP groups resulted in cell defects that were observed in more than one embryo, such as defects in cell cycle length, cell division timing, lineage-specific gene expression, cell division angle, and relative cell position (Figure 1B, iii; Supplemental Table S3). For TF and all protein-coding genes, depletion of 25.0% and 24.5% of genes, respectively, resulted in cell defects. Another study using RNAi and direct cell lineage tracing assigned 201 essential genes with roles in regulating lineage differentiation (Du et al. 2015); these include 27 RBP genes and 15 TF genes (Supplemental Table S3).

Finally, we also examined the conservation of RBPs. Among the RBP groups, 62.1-86.2% had a 1:1 ortholog with *C. briggsae* (Figure 1B, iv; Supplemental Table S1). On the other hand, TF and all protein-coding genes had 58.8% and 55.5% 1:1 orthologs in *C. briggsae*, respectively (Large et al. 2024). We similarly examined predicted human orthologs using OrthoList 2 (Kim et al. 2018), with all RBP and TF genes having significantly higher conservation compared to all protein-coding genes (Figure 1B, v; Supplemental Table S1).

### Many RBP genes have cell type-specific expression

While RBPs are known to be enriched for germline expression (Wang et al. 2009; Albarqi and Ryder 2023), it is not known whether they are enriched for cell type-specific expression overall or typically have broader expression. We asked whether RBP genes have broader or more cell type-specific expression than other genes using embryonic single-cell RNA-sequencing (scRNA-seq) data (Packer et al. 2019) and the *Tau* metric of expression specificity (Yanai et al. 2005; Large et al. 2024). *Tau* ranges from 0 to 1, with larger values corresponding to expression restricted to fewer cell types and lower values corresponding to broad expression. Group 1-3 RBP genes displayed bimodal distributions of *Tau*, with a bias toward broad expression or similar numbers having broad vs cell type-specific expression (Figure 2A-2C, 2E; Supplemental Table S1). This contrasted with TF genes, of which only 16.4% had broad expression (*Tau* < 0.4), similar to the rate seen in the full proteome (Figure 2E-2F; Supplemental Table S1). In line with their housekeeping functions, Group 4 RBP genes, which include many ribosomal protein genes, displayed the least cell type-specificity, with 75.1% of genes having broad expression (Figure 2D; Supplemental Table S1).

**Figure 2.**
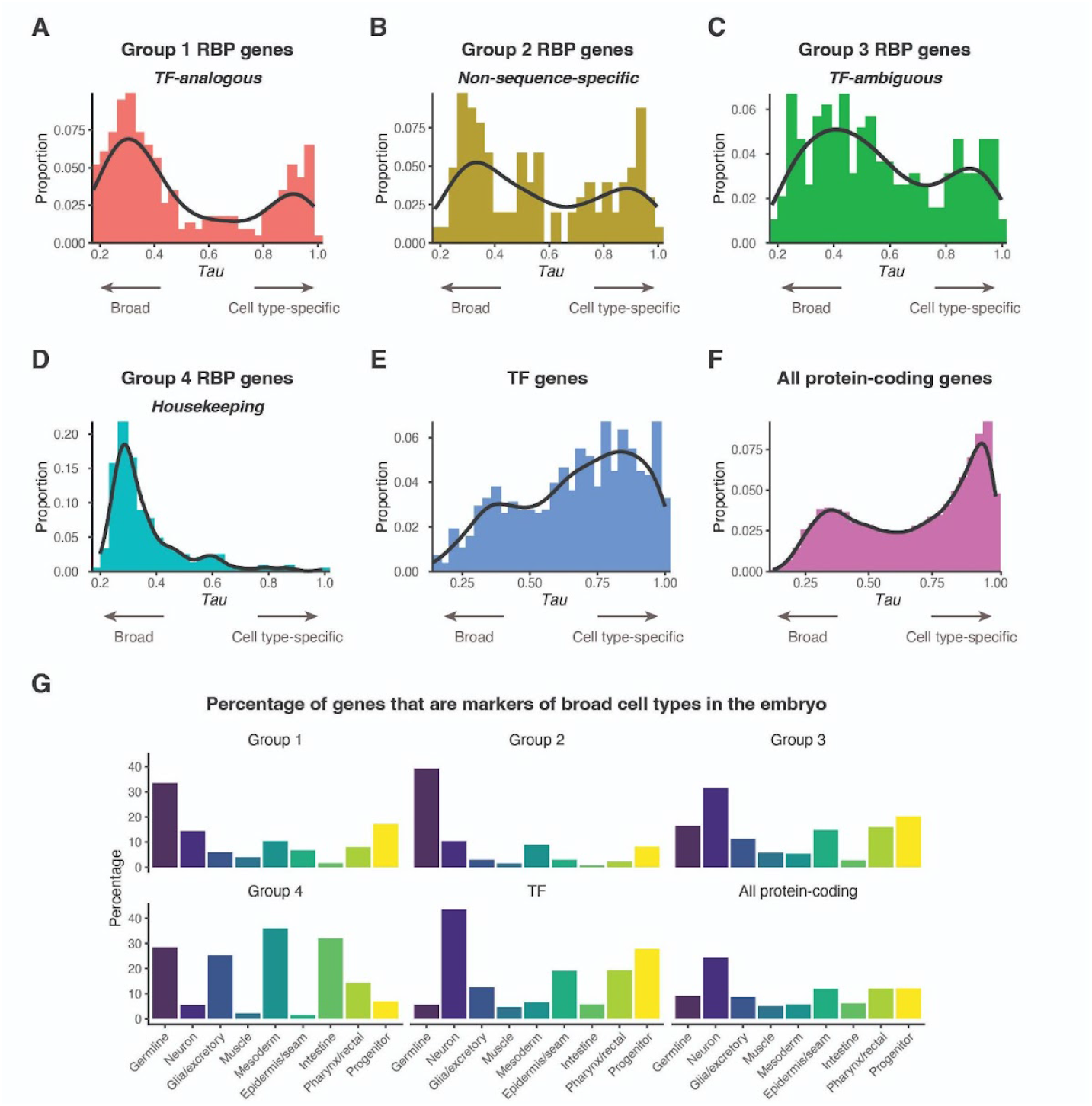
RBP genes have cell type-specific expression in the embryo. *Tau* distribution of (A) Group 1 RBP genes, (B) Group 2 RBP genes, (C) Group 3 RBP genes, (D) Group 4 RBP genes, (E) TF genes, and (F) all protein-coding genes. *Tau* was only included for genes with an expression of at least 5 transcript per million at any stage from a whole-embryo RNA-sequencing time series (Hashimshony et al. 2015). (G) Percentage of genes from each category that were markers of different broad cell types in the *C. elegans* embryo.

To further explore the cell type-specific expression of the RBP genes, we examined which of them were identified as cell type markers in an embryo scRNA-seq data (Figure 2G; Supplemental Table S1) (Large et al. 2024). Markers were defined as genes with significantly higher expression in a given cell type compared to others (> 1.5-fold, adjusted P < 0.05; see *Materials and Methods* for details). By these criteria, 481 TF genes (68.7%) were markers of at least one terminal or progenitor embryonic cell type. The terminal cell types with the highest amount of TF gene markers were neuronal, pharynx/rectal, and epidermis/seam cells (43.4%, 19.3%, and 19.1%, respectively). In contrast, a lower proportion of Group 1 RBP genes (139, 55.4%), are markers. The terminal cell type with the highest amount of Group 1 RBP gene markers was the germline (33.5%), consistent with past studies showing RBPs enriched in the germline (Wang et al. 2009; Albarqi and Ryder 2023). However, many other terminal cell types, including neurons and mesoderm (14.3% and 10.4%, respectively), had multiple group 1 RBP gene markers, providing candidates for post-transcriptional regulators of these cell types. Similar marker trends were observed for Group 2 RBP genes, while Group 3 genes (many of which could encode either RBPs or TFs) had similar patterns to TFs.

### Many RBP genes have developmental stage-specific expression

We assessed the patterns of RBP and TF gene expression across developmental stages of embryogenesis. We categorized their expression over time in a whole-embryo RNA-sequencing time series (Hashimshony et al. 2015) into eight clusters (Figure 3; Supplemental Table S2). Six of these clusters exhibited peaks in expression, suggesting stage-enriched expression that spanned the range of embryonic development. We refer to them here based on their time of maximum expression (e.g. “Maternal 1”, “Early 2”, etc). TF genes were most likely to be expressed in the “Middle” and “Late” clusters (Figure 3C-3D). In contrast, Group 1-3 RBP genes were most enriched in the “Maternal” and “Early” clusters, indicating more frequent roles earlier in embryogenesis (Figure 3A-3B).

**Figure 3.**
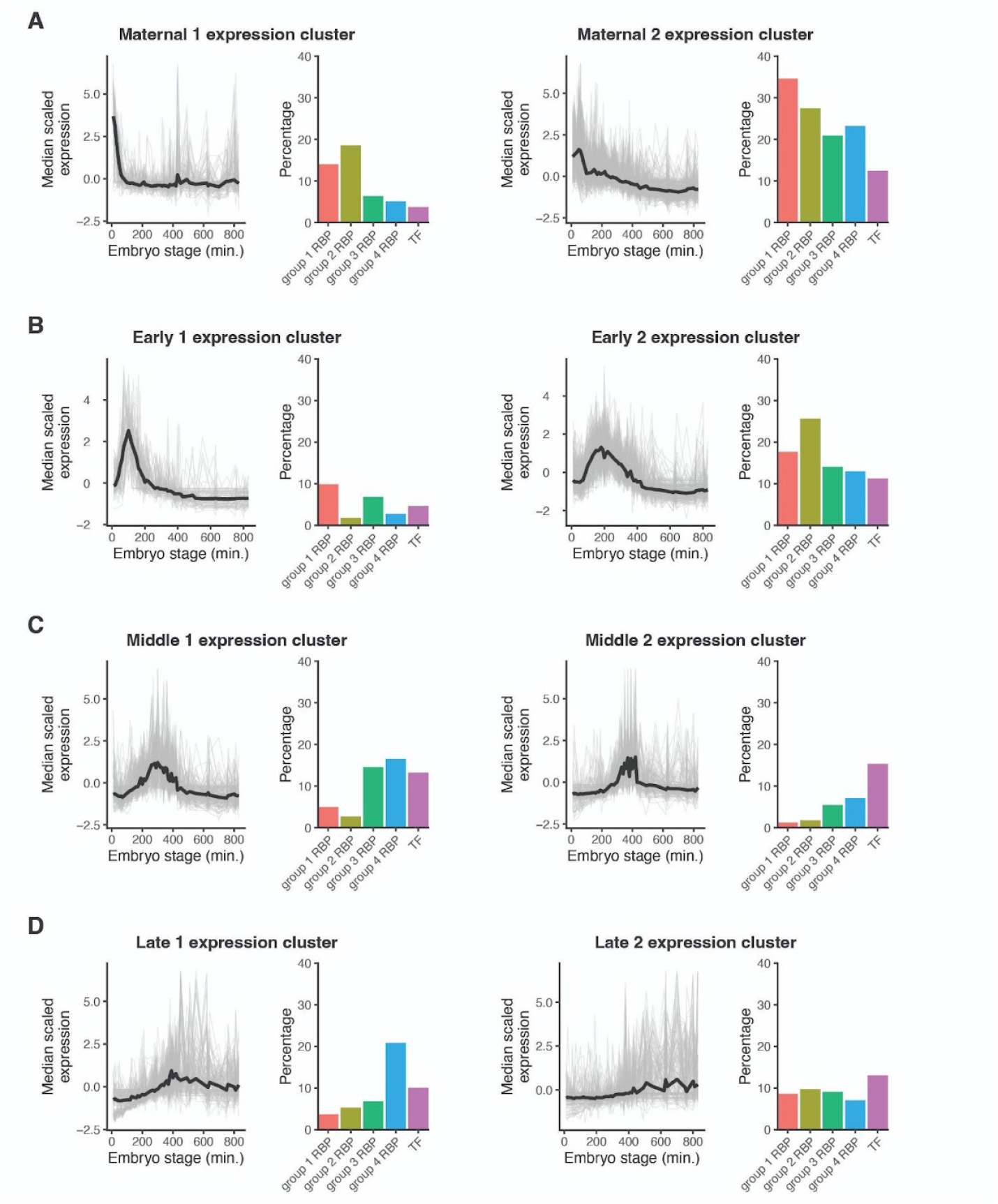
RBP genes have developmental stage-specific expression in the embryo. (A-D) The percentages of RBP and TF genes that fell into specific clusters of expression throughout embryogenesis. Expression data for RBP and TF genes was taken from a whole-embryo RNA-sequencing time series (Hashimshony et al. 2015) and hierarchically clustered.

There were also broad differences in expression across the life cycle of *C. elegans*. We examined the expression over time of each RBP and TF gene group in a time-resolved transcriptome of *C. elegans* embryonic and post-embryonic stages (Boeck et al. 2016) (Figure 4; Supplemental Table S4). Across all categories, the majority of RBP and TF genes appeared to be most highly expressed in the embryo. However there was diversity among RBP and TF genes for expression across the larval and young adult stages. For example, there were many examples where gene expression increased from L1 to the young adult stage for Group 1 RBP genes, while this was less common for TF genes. Together, these analyses indicate that many RBP genes exhibit specificity in their expression across cell types and stages, potentially allowing for context-specific regulation of their RNA targets.

**Figure 4.**
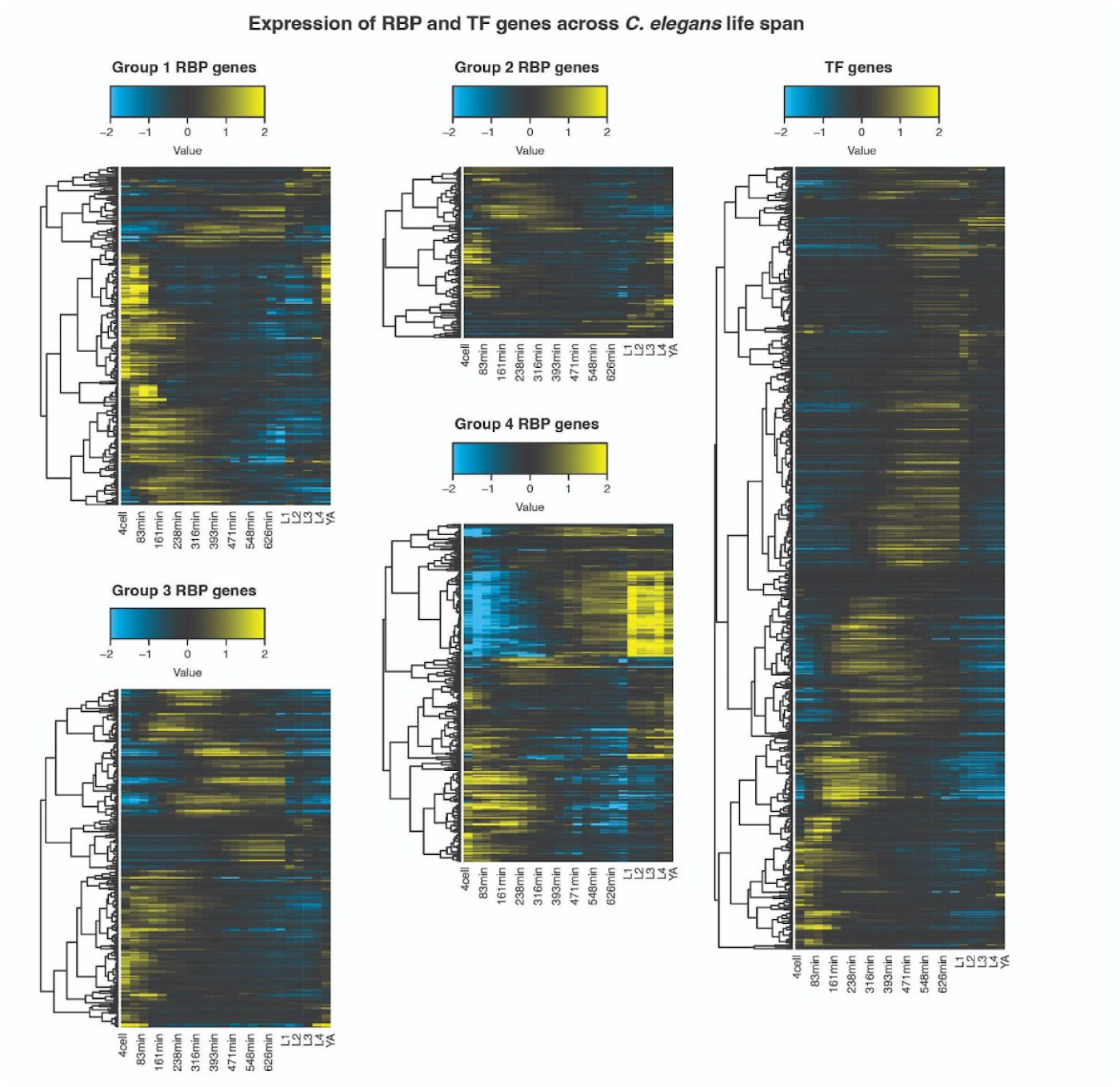
RBP genes have developmental stage-specific expression across the life cycle of *C. elegans*. Heatmaps of gene expression across embryonic, larval, and young adult stages in *C. elegans* across different categories of genes. Embryonic stages on the left half of the plots are given in minutes, post-two-cell embryo. Expression data was taken from a time-resolved transcriptome of *C. elegans* across embryonic and post-embryonic stages (Boeck et al. 2016). Expression was placed on a log_2_ scale and mean-centered by row in the heatmaps. Genes were ordered based on hierarchical clustering.

### Many Group 1 RBP genes are enriched in the germline

We focused our remaining analyses on Group 1 RBPs, as these RBPs bind targets via RNA motifs in a way most analogous to TFs. As described above, many RBP genes are markers of embryonic germline precursor cells (GPCs), in addition to their known enrichment in the adult germline (Wang et al. 2009; Albarqi and Ryder 2023). The embryonic GPCs are transcriptionally quiescent, potentially requiring the use of RBPs for post-transcriptional regulation. In addition, early- vs late-embryo GPCs differ in their transcriptome in part due to differential RNA degradation (Peng et al. 2024). We compared RBP gene expression in the GPCs across time using previously defined pseudotime bins in a scRNA-seq atlas (Packer et al. 2019). We found 49 RBP genes increase in relative expression from the first pseudotime bin to the last, while 35 decrease in expression (Figure 5). Several of the genes with the greatest fold-changes in expression over time encode proteins with well-known roles in the *C. elegans* germline. For example, the CCCH domain-containing protein POS-1 and the RRM domain-containing protein SPN-4 can interact with one another and have known early roles in regulating the translation of maternal mRNAs (Ogura et al. 2003); *pos-1* and *spn-4* had strong decreases in expression over time in the germline (Figure 5A). On the other hand, Germ Line DEAD-box Helicase (GLH) family are CCHC domain-containing proteins that are components of P granules and have roles in the regulation of germline RNAs (Gruidl et al. 1996; Beshore et al. 2011); *glh-1/2/3/4* had strong increases in relative expression over time in the germline (Figure 5B). The proteins encoded by these early and late-enriched RBP genes in the embryonic germline provide candidate regulators of GPC maturation.

**Figure 5.**
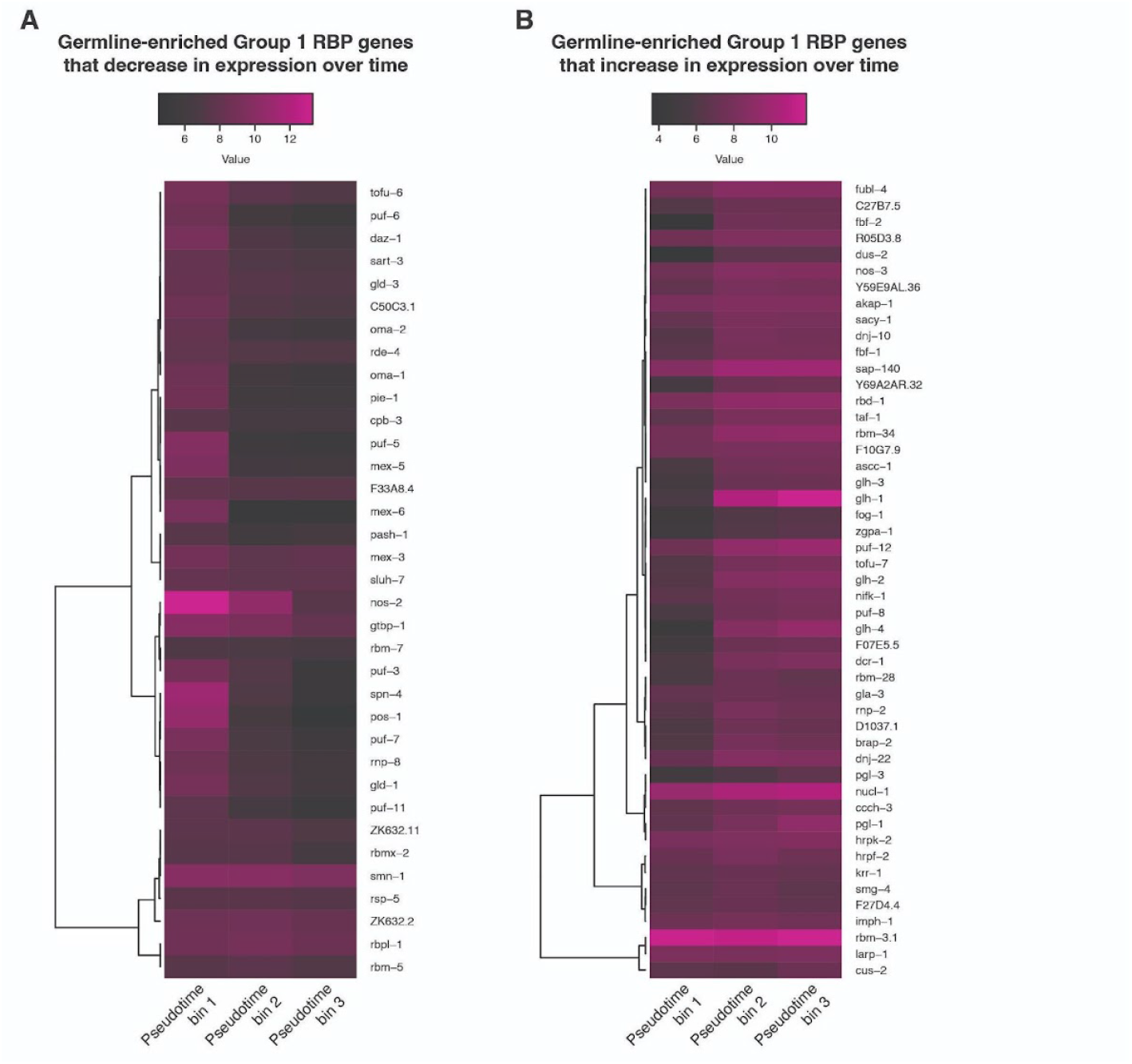
Many Group 1 RBP genes are enriched in the germline. (A) Heatmap of the germline-specific expression of germline-enriched Group 1 RBP genes that decrease in expression over pseudotime. Expression data was taken from a *C. elegans* embryo single-cell atlas (Packer et al. 2019) and placed on a log_2_ scale. Genes were ordered based on hierarchical clustering. (B) Heatmap of the germline-specific expression of germline-enriched Group 1 RBP genes that increase in expression over pseudotime. Expression data was taken from a *C. elegans* embryo single-cell atlas (Packer et al. 2019) and placed on a log_2_ scale. Genes were ordered based on hierarchical clustering.

### Many zygotic-only Group 1 RBP genes are terminal cell type markers

The vast majority of the germline-enriched RBP genes are maternally expressed; to understand the remaining zygotic RBPs, we examined the Group 1 RBP genes not detected in the zygote (see *Materials and Methods*). Of the 60 zygotic-only Group 1 RBP genes, 23 were markers of at least one terminal cell type, with most being markers for specific neuronal cell types (Figure 6A; Supplemental Table S5). For example, *cpb-2*, a homolog of cytoplasmic polyadenylation element binding genes (Luitjens et al. 2000), has enriched expression in the ADF sensory neuron. Other Group 1 RBP genes have enriched expression in many neuronal cell types, such as *unc-75*, which encodes a protein that is part of the CELF family of RBPs and is involved in neuron-specific alternative splicing (Kuroyanagi et al. 2013). Expression of *unc-75* is enriched in 26 terminal cell types that span both ciliated and non-ciliated neurons.

**Figure 6.**
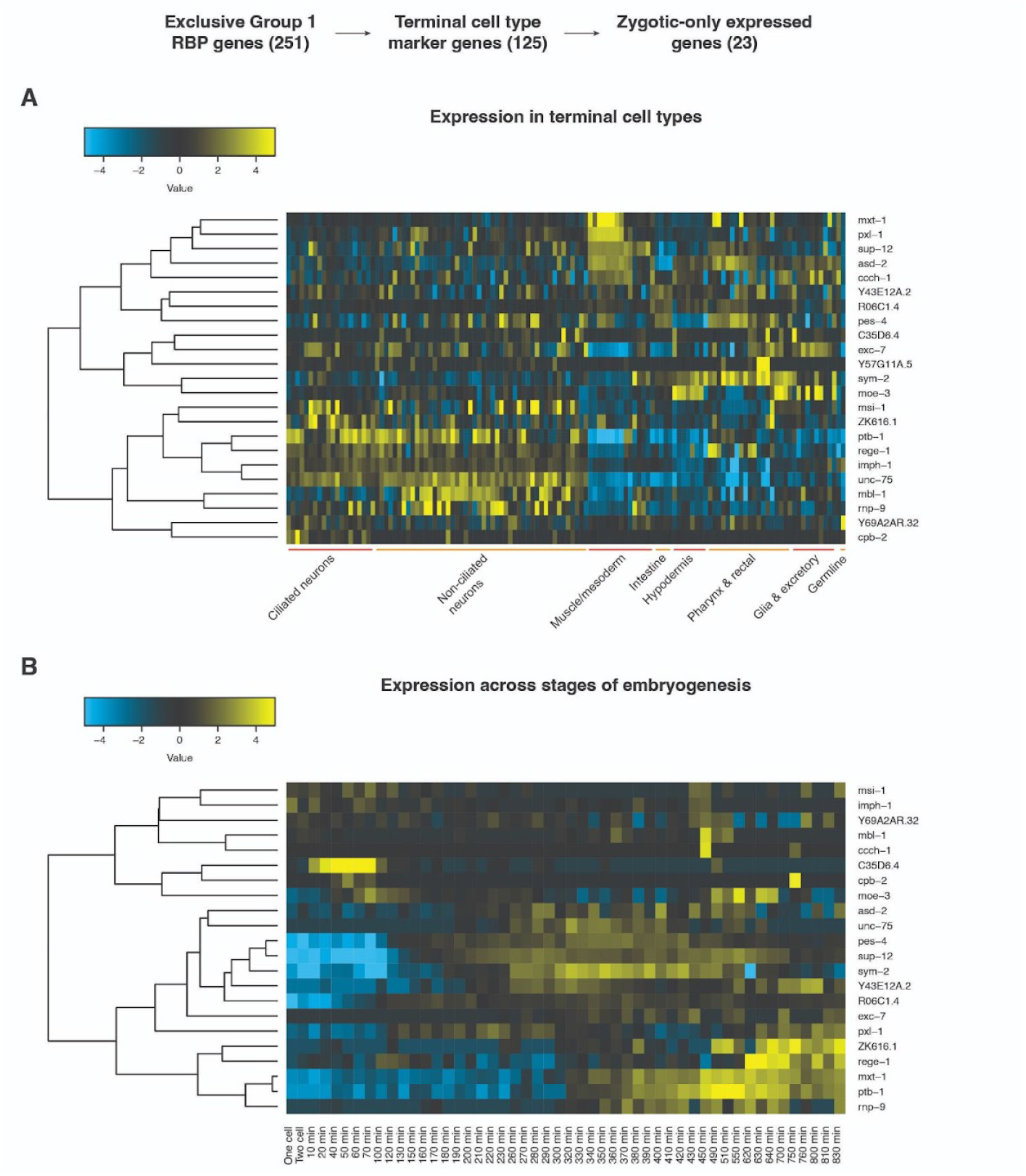
Many zygotic-only Group 1 RBP genes are terminal cell type markers. Group 1 RBPs were examined if they were characterized as a terminal cell type marker and as only zygotically expressed. (A) Heatmap of the expression of such genes in broad terminal cell types. Expression data was taken from a *C. elegans* embryo single-cell atlas (Packer et al. 2019), placed on a log_2_ scale, and mean-centered by row in the heatmap. Genes were ordered based on hierarchical clustering. (B) Heatmap of the expression of such genes across stages of embryogenesis. Expression data was taken from a whole-embryo RNA-sequencing time series (Hashimshony et al. 2015), placed on a log_2_ scale, and mean-centered by row in the heatmap. Genes were ordered based on hierarchical clustering.

Many zygotic-only Group 1 RBP genes were also enriched in muscle/mesodermal cells. For example, *pxl-1*, a paxillin homolog that encodes a protein required for muscle contraction in the pharynx (Warner et al. 2011), has enriched expression in body wall muscle. Another gene, *asd-2*, has enriched expression in body wall muscle, pharyngeal muscle, and the neurons AVL, URB, and URA; ASD-2 is involved in the regulation of alternative splicing (Ohno et al. 2008; Ohno et al. 2012). Several of these zygotic-only Group 1 RBP genes with cell type-specific expression also displayed dynamic expression across developmental stages (Figure 6B; Supplemental Table S5), highlighting the potential for multidimensional context-specific RNA regulation. For instance, *unc-75* and *asd-2* were both part of clusters from Figure 3 that peaked in expression around the time that terminal differentiation begins in the embryo.

### Group 1 RBP genes have distinct mRNA decay dynamics

wRBP1.0 identified the likely extensive transcriptional and post-transcriptional regulation of RBPs (Tamburino et al. 2013). The resource found that more TFs bind to typical RBP promoters than to promoters of other genes. RBP proteins were also found to bind more frequently to RBP-encoding mRNAs than to other mRNAs, and RBP genes were more often alternatively spliced compared to other genes. To expand upon these observations, we examined the mRNA decay dynamics of Group 1 RBP genes using an atlas of mRNA decay rates in the *C. elegans* embryo (Peng et al. 2024).

Among genes with measurable mRNA half-lives in the embryo, we found that Group 1 RBP transcripts had longer half-lives overall compared to TF and other protein-coding genes, with a median half-life of 45 minutes (Figure 7A; Supplemental Table S2). When breaking down the Group 1 RBP genes by which RBDs they encoded, there were small differences in their half-life distributions, though none reached statistical significance (Figure 7B). Group 1 RBP genes encoding the CCCH, RRM, and KH domains displayed a wide range of median mRNA half-lives (38, 45, and 55 minutes, respectively). As mRNA half-lives in the *C. elegans* embryo were found to be shorter overall in earlier stages of embryogenesis (Peng et al. 2024), we examined whether there may be differences in expression timing of genes encoding different RBDs. The median scaled expression of Group 1 RBP genes encoding CCCH domains peaked before that of Group 1 RBP genes encoding RRM or KH domains (Figure 7C). In addition, the expression of genes encoding KH domains peaked and plateaued in expression relatively late in embryogenesis. This suggests specific families of RBPs may act preferentially at different stages of development.

**Figure 7.**
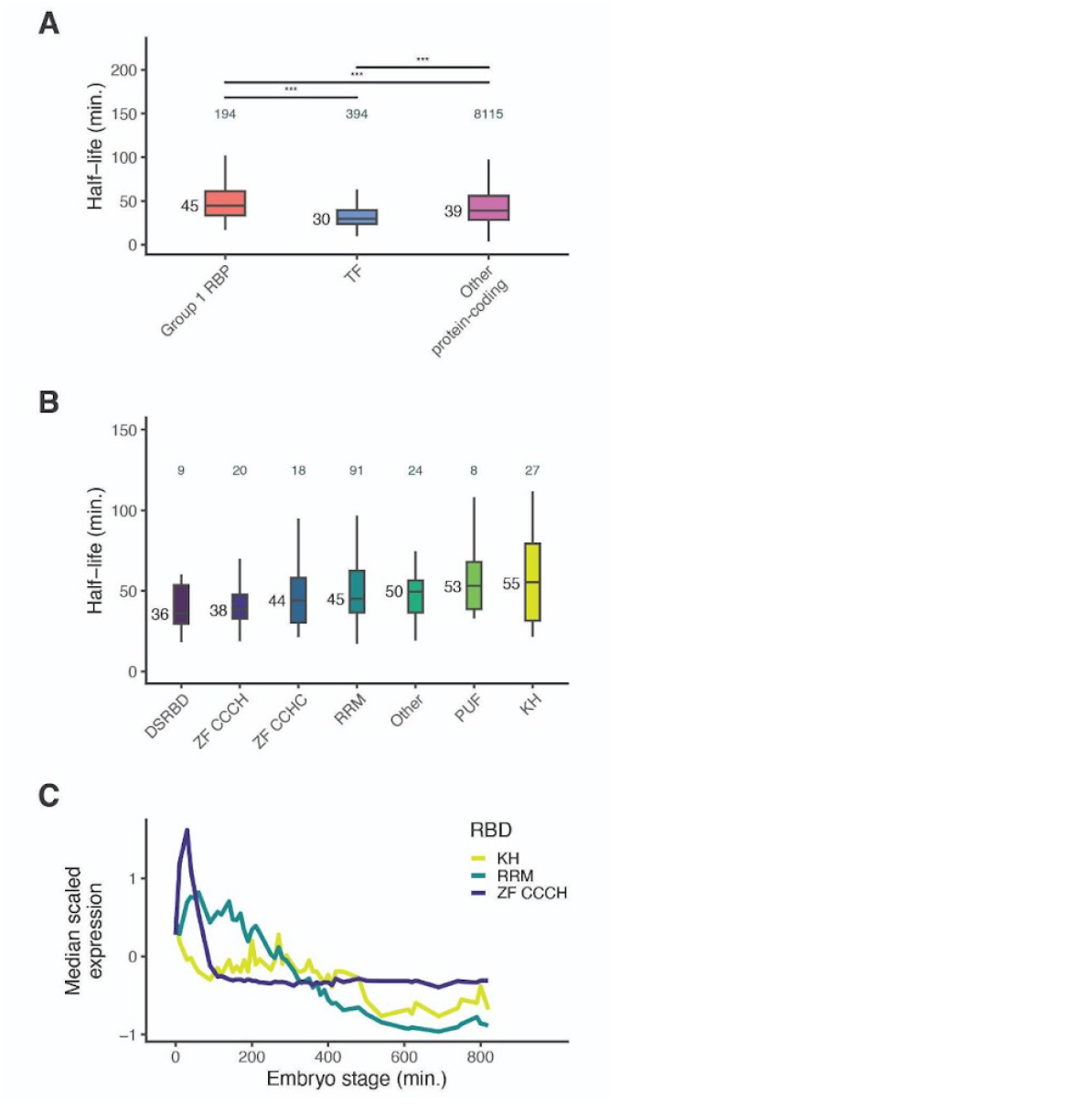
Group 1 RBP genes have distinct mRNA decay dynamics. (A) Box plots showing the pseudobulk mRNA half-life distributions of genes within each category. Half-life data was taken from a *C. elegans* embryo atlas of mRNA decay rates (Peng et al. 2024). Numbers to the left of the box plots are median half-lives within each group. Numbers above box plots are the number of genes within each group. P-value comparing median half-lives was calculated using the Wilcoxon rank sum test. (B) Box plots showing the pseudobulk mRNA half-life distributions of genes within each RBD category. Half-life data was taken from a *C. elegans* embryo atlas of mRNA decay rates (Peng et al. 2024). Numbers to the left of the box plots are median half-lives within each group. Numbers above box plots are the number of genes within each group. P-value comparing median half-lives was calculated using the Wilcoxon rank sum test. (C) Median scaled expression of gene subsets using data from a whole-embryo RNA-sequencing time series (Hashimshony et al. 2015).

## DISCUSSION

Characterization of RBP regulatory networks throughout development has yet to be carried out as comprehensively as it has been for TF regulatory networks. In this work, we expand upon an existing RBP compendium in *C. elegans* that highlights the potential for RBPs to be extensively regulated at multiple levels (Tamburino et al. 2013). We have updated the list of predicted RBPs to 928 proteins (254 Group 1 proteins, 139 Group 2 proteins, 257 Group 3 proteins, and 278 Group 4 proteins). In addition, we explore the potential for RBPs to regulate developmental gene expression patterns. This was done by examining RBP gene expression throughout *C. elegans* embryogenesis and the developmental defects that arise when RBPs are depleted.

### RNA-binding proteins have important roles in appropriate C. elegans development

We found that RBP genes had substantial developmental defects associated with their depletion in *C. elegans* (Figure 1B). These defects ranged from embryonic lethality to cell division defects in the embryo, suggesting that many RBP genes play key roles in ensuring proper embryonic development. We also found in general that RBP genes in *C. elegans* had higher percentages of 1:1 orthologs with *C. briggsae* compared to TF and all protein-coding genes (Figure 1B, iv). This supports the idea that RBPs are particularly well-conserved as a result of their developmental importance, and studies of RBPs in *C. elegans* may be broadly generalizable. Future work could determine whether the phenotypes and conservation of gene-specific Group 1 RBPs compared to TFs result from their roles in more cell types, as evidenced in their trend towards broader expression, or the number or nature of their target genes.

### Group 1 RNA-binding proteins have the potential to regulate gene expression in a highly specific manner

A key step toward characterizing RBP regulatory networks throughout development is to determine where and when RBPs are expressed. To this end, we examined the expression of RBP genes throughout *C. elegans* embryogenesis. While many Group 1 RBP genes displayed greater expression early on in embryogenesis as well as more widespread expression across cell types compared to TF genes (Figure 2; Figure 3), there were Group 1 RBP genes that peaked at all embryonic stages, indicating that they could regulate their RNA targets in a highly specific manner.

The majority of Group 1 RBP genes that were enriched in a terminal cell type were enriched in the germline and maternally expressed. During germline and early embryonic development, contexts in which cells are largely transcriptionally quiescent, post-transcriptional mechanisms that regulate gene expression become more predominant (Lee and Schedl 2006 Apr 18). Consequently, much of the more well-characterized RBPs in *C. elegans* have functions during these periods of development, particularly as regulators of translation (Lee and Schedl 2006 Apr 18).

Nevertheless, post-transcriptional regulation of gene expression is also important in somatic development, and several RBPs have somatic tissue-specific defects upon their depletion (Lee and Schedl 2006 Apr 18). We identified 23 zygotic-only group 1 RBP genes, 21 of which were markers of somatic terminal cell types (Figure 6A). Several of these RBPs, such as EXC-7 and UNC-75, are involved in regulating alternative mRNA splicing (Kuroyanagi et al. 2013; Tan and Fraser 2017). Several more, such as MBL-1, PTB-1, and Y57G11A.5, are predicted to be involved in RNA splicing as well (Lee et al. 2018). While many germline- and soma-enriched Group 1 RBPs remain to be characterized in detail, these findings suggest that many somatic tissue-enriched Group 1 RBPs may function in RNA processing events such as splicing rather than in post-transcriptional events taking place in the cytoplasm.

#### Expression of genes encoding Group 1 RNA-binding proteins is extensively regulated

The previous wRBP1.0 compendium demonstrated the potential for extensive regulation of RBP genes and the proteins themselves (Tamburino et al. 2013). We expanded upon the study of RBP gene regulation by examining the mRNA half-lives of Group 1 RBP genes using an atlas of mRNA decay rates in the *C. elegans* embryo (Peng et al. 2024). We observed differential mRNA decay dynamics between Group 1, TF, and other protein-coding genes, with Group 1 RBP genes having longer mRNA half-lives overall (Figure 5A). Differential decay dynamics may also exist between Group 1 RBP transcripts encoding different RBDs. Though none of the differences in median half-lives reached statistical significance, transcripts encoding the CCCH domain appeared to have relatively rapid decay compared to transcripts encoding RRM or KH domains (Figure 7B). While this may be due in part to differences in expression timing of these genes, it also appears likely that CCCH-encoding transcripts are specifically targeted for rapid degradation due to their overall rapid decreases in expression in the early embryo (Figure 5C). MEX-5 and MEX-6 are nearly identical CCCH finger proteins that have roles in polarity establishment in the early *C. elegans* embryo (Schubert et al. 2000). We previously identified differential regulation of *mex-5* and *mex-6* transcript turnover, with faster decay rates observed in the germline compared to in the soma (Peng et al. 2024). While the mechanism behind this remains to be elucidated, this highlights the potential for extensive regulation of the degradation of CCCH-encoding transcripts. Overall, our findings update the previous wRBP1.0 compendium and further emphasize the complexity of RBP regulatory networks in early development.

## MATERIALS AND METHODS

### Identification of predicted RBPs

InterPro protein domains were retrieved from the WormBase ParaSite database (Howe et al. 2017) on September 23, 2024. The proteome was then searched for each of 17 RBDs that were included previously in wRBP1.0: RRM, KH, PUF, CCCH, CCHC, DSRBD, RGG Box, La, Helicase, PAZ, PIWI, Argonautes, NTF2, C2H2, SAM, Cold shock, and Sm/Lsm (Tamburino et al. 2013). As with wRBP1.0, domains were filtered based on whether they were identified in any of the following databases: Pfam, SMART, Superfamily, or ProSite. Gene names from wRBP1.0 were updated to align with the *C. elegans* genome build WS290, and dead genes or those that are transposon in origin were removed from the compendium.

### Defining RBP, TF, and other protein-coding genes

Putative TF genes were taken from the compendium, wTF3.0 (Fuxman Bass et al. 2016). A list of protein-coding genes was retrieved from the WormBase ParaSite database (Howe et al. 2017) on September 23, 2024. For downstream comparative analyses, we limited Group 1 RBPs to those mutually exclusive with TFs; however, proteins predicted to potentially be both RBPs and TFs were categorized as RBPs if they contained a CCCH domain. Group 2 RBPs were excluded of TFs. As Group 3 RBPs are by definition ambiguous in having RNA- or DNA-binding functions, putative TFs were kept in this group for downstream analyses. Putative RBPs from all four groups were excluded from TFs analyzed.

### Defining maternal and zygotic-only genes

Zygotic-only genes were determined using a whole-embryo time series RNA-sequencing dataset of the *C. elegans* embryo (Hashimshony et al. 2015) and a single-cell RNA-sequencing dataset on each cell of the *C. elegans* embryo through the 16-cell stage (Tintori et al. 2016). Genes with transcript per million (TPM) >= 10 within one-cell embryos in the whole-embryo dataset or an average reads per kilobase million (RPKM) >= 20 within one-cell embryos in the single-cell dataset were considered to be maternally-expressed genes. All other genes were considered to be zygotic-only genes.

### Characterizing expression of RBP and TF genes across embryonic development

Gene expression patterns for RBP and TF genes from a whole-embryo time series RNA-sequencing dataset of the *C. elegans* embryo (Hashimshony et al. 2015) were first scaled using the ‘scale’ function in R. The similarity in expression patterns between pairs of genes was determined using the Pearson correlation coefficient, and the correlations were turned into distance measures by subtracting them from 1 and passing them to the ‘as.dist’ function. Hierarchical clustering was performed on these distance values using the ‘hclust’ function. A cluster number of 10 was chosen based on visualization with a dendrogram, with the two clusters with the lowest amount of genes removed for clarity in downstream analyses.

A heatmap for zygotic-only Group 1 RBP genes that are terminal cell type markers was also created using gene expression data from the whole-embryo time series RNA-sequencing dataset of the *C. elegans* embryo (Hashimshony et al. 2015). A pseudocount of 1 was applied to the transcripts per million (TPM) of each gene at each time point, then the log_2_ was taken of these modified expression levels. Hierarchical clustering of this gene expression was performed similarly as above, and a heatmap visualization was generated using the ‘heatmap.2’ function.

### Characterizing expression of RBP genes across embryonic cell types

Terminal cell type marker genes in the *C. elegans* embryo were determined as in the single-cell transcriptome atlases of *C. elegans* and *C. briggsae* (Large et al. 2024), though a more stringent log_2_ fold-change cutoff of 1.5 was used instead. Group 1 RBP genes that were enriched in the germline were further examined using germline-specific expression data from a *C. elegans* embryo single-cell atlas (Packer et al. 2019). This data was used to create heatmaps of expression for genes that increased or decreased in expression over time in the germline. A pseudocount of 1 was applied to the transcripts per million (TPM) of each gene within each pseudotime bin, then the log_2_ was taken of these modified expression levels. Hierarchical clustering of this gene expression was performed similarly as in the above section, and a heatmap visualization was generated using the ‘heatmap.2’ function.

## Data availability

R codes associated with the updated list of *C. elegans* RNA-binding protein genes and downstream comparative analyses are available at GitHub (https://github.com/fe-peng/celegans_RBP_compendium).

## Acknowledgements

We thank members of the Murray lab for providing valuable discussion and comments on the manuscript. This work was funded by R35GM153497 and F31HD10785.

